# Craniofacial diversity across Danionins and the effects of TH status on craniofacial morphology of two Danio species

**DOI:** 10.1101/2023.08.09.552728

**Authors:** Stacy Nguyen, Rachel S Lee, Emma Mohlmann, Gabriella Petrullo, John Blythe, Isabella Ranieri, Sarah McMenamin

**Affiliations:** Biology Department, Boston College, 140 Commonwealth Avenue, Chestnut Hill, MA 02467

## Abstract

The model zebrafish (*Danio rerio*) belongs to the Danioninae subfamily with a range of informative phenotypes. However, the craniofacial diversity across the subfamily is not fully described. To better understand craniofacial phenotypes across Danioninae we used microCT and 3D geometric morphometrics to capture skull shapes from nine species. The *Danio* species examined showed largely similar skull shapes, although *D. aesculapii*, the sister species to *D. rerio* showed a unique morphology. Two non-*Danio* species examined, *Chela dadiburjori* and *Devario aequipinnatus* showed distinct skull morphologies unique from those of other species examined. Thyroid hormone regulates skeletal development and remodeling, and we asked if changes in developmental thyroid hormone metabolism could underlie some of the craniofacial diversity across Danioninae. We reared two *Danio* species under altered thyroid profiles, finding that hypothyroid individuals from both species showed corresponding morphological shifts in skull shape. Hypothyroid *Danios* showed skull morphologies closer to that of *Chela* and unlike any of the examined wild-type *Danio* species. We provide an examination of the evolved craniofacial diversity across Danioninae, and demonstrate that alterations to thyroid hormone have the capacity to create unique skull phenotypes.

## Introduction

Zebrafish (*Danio rerio*) are a powerful model in developmental biology and biomedicine, and have emerged as a tractable system for studying vertebrate skeletogenesis and craniofacial development (e.g. see 1, 2, 3). There is growing interest in the natural phenotypic diversity across the broader *Danio* genus and more distantly related relatives of *D. rerio* (e.g. see 4, 5, 6). Placing *D. rerio* into this larger phylogenetic context can help provide ecological, evolutionary, and functional depth for zebrafish research, can suggest avenues for ongoing molecular and biomedical efforts, and can identify informative phenotypes for genomic and developmental analysis.

Belonging to the Cypriniform order, the Danioninae subfamily contains at least 20 genera and over 100 identified species (7, 8). Several of these species are common in India and southeast Asia, and are endemic to a number of major hydrological basins (9). Danioninae are typically flexible omnivores, generally consuming insects, crustaceans, algae and detritus (10). Species show a range of morphological characteristics adapted to their slightly different ecologies (11, 12). Although the systematics of Danioninae have been contentious, modern genomic methods have increasingly arrived on consensus relationships across the group (7, 13). The subfamily contains three major genera: *Chela, Devario* and *Danio* (8, 14).

Many of the characteristics making *D. rerio* an outstanding research model—small size, ease of care and breeding—make other Danioninae attractive as both aquarium fish for hobbyists and as a research subjects for biologists (e.g. see 15, 16). The diversity that exists within the Danioninae subfamily can be leveraged to ask powerful questions about mechanisms of evolutionary divergence. Different *Danio* species exhibit spectacularly diverse pigmentation, and the evolution of these pigment patterns is being studied at developmental and molecular levels (see 17, 18).

Although not as conspicuous as diversity of pigment patterns, elements of the craniofacial skeleton differ between Danioninae species (4, 19). Skull diversity across Danioninae could form the basis for fruitful evolutionary comparisons, laying groundwork for comparative molecular work. To this end, here we use microcomputed tomography (microCT) examine the overall craniofacial diversity across several Danioninae, including seven *Danio* species as well as a *Devario* and *Chela*.

Thyroid hormone (TH) regulates skeletal development, and in zebrafish and other Cypriniformes, the hormone is known to specifically shape elements of the craniofacial skeleton (19-21). Here, we asked if some of the overall craniofacial differences we observed across Danioninae mirrored shifts associated with changes to TH metabolism. To test this, we asked whether *Danio* species reared under altered TH profiles showed craniofacial morphology that more closely resembled that of other Danioninae species. Our results both provide insights into the natural diversity in Danioninae head shape and suggest that alterations to TH have the capacity to create unique overall craniofacial phenotypes.

## Methods

### *Danio* lines and Danioninae species

All experiments were performed using an approved protocol in accordance with the Institutional Animal Care and Use Committee at Boston College. *D. rerio* and *D. albolineatus* were from lines transgenic for *tg:nVenus-2a-nfn* (22); below details the thyroid ablation procedure used to produce hypothyroid (HypoTH) individuals. Hyperthyroid (HyperTH) *D. rerio* were of the mutant line *opallus* (22). *D. nigrofasciatus* and *D. aesculapii* were obtained from the Parichy Lab (University of Virginia) and were reared at Boston College. All other Danioninae species were identified by retailers and purchased from tropical fish suppliers: *Devario aequipinnatus*, Petco, Shrewsbury, MA; *C. dadiburjori* and *D. erythromicron*: Uncle Ned’s Fish Factory, Millis, MA; *D. kyathit* and *D. margaritatus*: Lucky’s Aquarium, Worcester, MA.

### Thyroid ablation

HypoTH *D. rerio* and *D. albolineatus* were produced, briefly, by treating 4 day-old larvae transgenic for *tg:nVenus-2a-nfn* with 10 mM metronidazole with 1% DMSO to induce thyroid ablation (see McMenamin et al, 2014). Control, wild-type fish were produced with the same thyroid ablation line treated only with 1% DMSO. All fish were reared under standard conditions (28°, 12:12 light cycles), recirculating water was carbon-filtered, and fish were fed TH-free food: rotifers fed with RotiGrow Plus (Reed Matriculture), Artemia, and pure Spirulina flakes (Pentair). HypoTH groups and their controls were fed a diet of artemia and spirulina flakes.

### Fish rearing

All experiments were performed using an approved protocol in accordance with the Institutional Animal Care and Use Committee at Boston College. *D. rerio, D. albolineatus, D. nigrofasciatus* and *D. aesculapii* were reared at 28°C on a 14:10 light:dark cycle and fed a diet of marine rotifers and adult pellet food flakes three times a day. Hypothyroid *D. rerio and D. albolineatus* were fed a diet of artemia and spirulina flakes.

### MicroCT scanning

Adult fish were euthanized in MS-222 and fixed in 4% paraformaldehyde for 24 hours. Standard length (SL) was measured from fixed specimens and SL was converted according to Parichy *et al* (23). Samples were placed between low-density foam molds and scanned with a SkyScan 1275 high resolution microCT system (Bruker, Kontich, Belgium) using a scanning resolution of 10.5 um and an x-ray source voltage of 45 kV and current of 200mA. Projection images were generated over 360° with a 0.2° rotation step and 4 averaging frames. Thresholding, ring artifact reduction, and beam hardening corrections were consistent across all scans during reconstruction using NRecon (Bruker, Kontich, Belgium).

### 3D modeling

3D models of the Danioninae heads were generated using the Segmentation Editor in Amira 6.5 (Thermo Fisher Scientific FEI, Hillsboro, Oregon, USA). The entire scan volume was loaded, and a pixel threshold was determined to differentiate bone from soft tissue. The lasso tool was used to select the corresponding pixels of the head region and added to a new material label.

The generated surface module was used to create a 3D model from the corresponding material label and the model was then exported as a .ply file. 3D models were further processed and simplified using MeshLab (24). Individual elements were segmented using the Segmentation Editor in Amira 6.5 (Thermo Fisher Scientific FEI, Hillsboro, Oregon, United States).

### Automatic landmarking

Four species (*D. rerio, D. aesculapii, D. kyathit*, and *D. nigrofasciatus*) were evaluated as potential templates, and *D. rerio* was determined to be the best template for automatic landmarking. Each 3D model was screened for quality, and specimens showing excessive gape or jaw mispositioning were excluded. A total of 3000 landmarks were automatically placed on each 3D model using the Automated Landmarking through the 3D slicer extension Pointcloud Alignment and Correspondence Analysis (ALPACA) (25, 26). General Procrustes alignment was performed to remove differences in positioning, size/scale, and orientation on the original dataset of landmark coordinates. A Principal Component Analysis (PCA) was performed using the SlicerMorph extension of 3D Slicer (27). Results were plotted onto a morphospace using the Geomorph package in RStudio (28).

## Results

### Danioninae exhibit a range of craniofacial shapes

Renderings of representative skulls from nine Danioninae species revealed differences in density and head shape (Fig 1). To quantitatively capture craniofacial shapes, we generated 3D models of the heads and automatically landmarked each head (Supplementary Fig 1).

**Figure 1.**
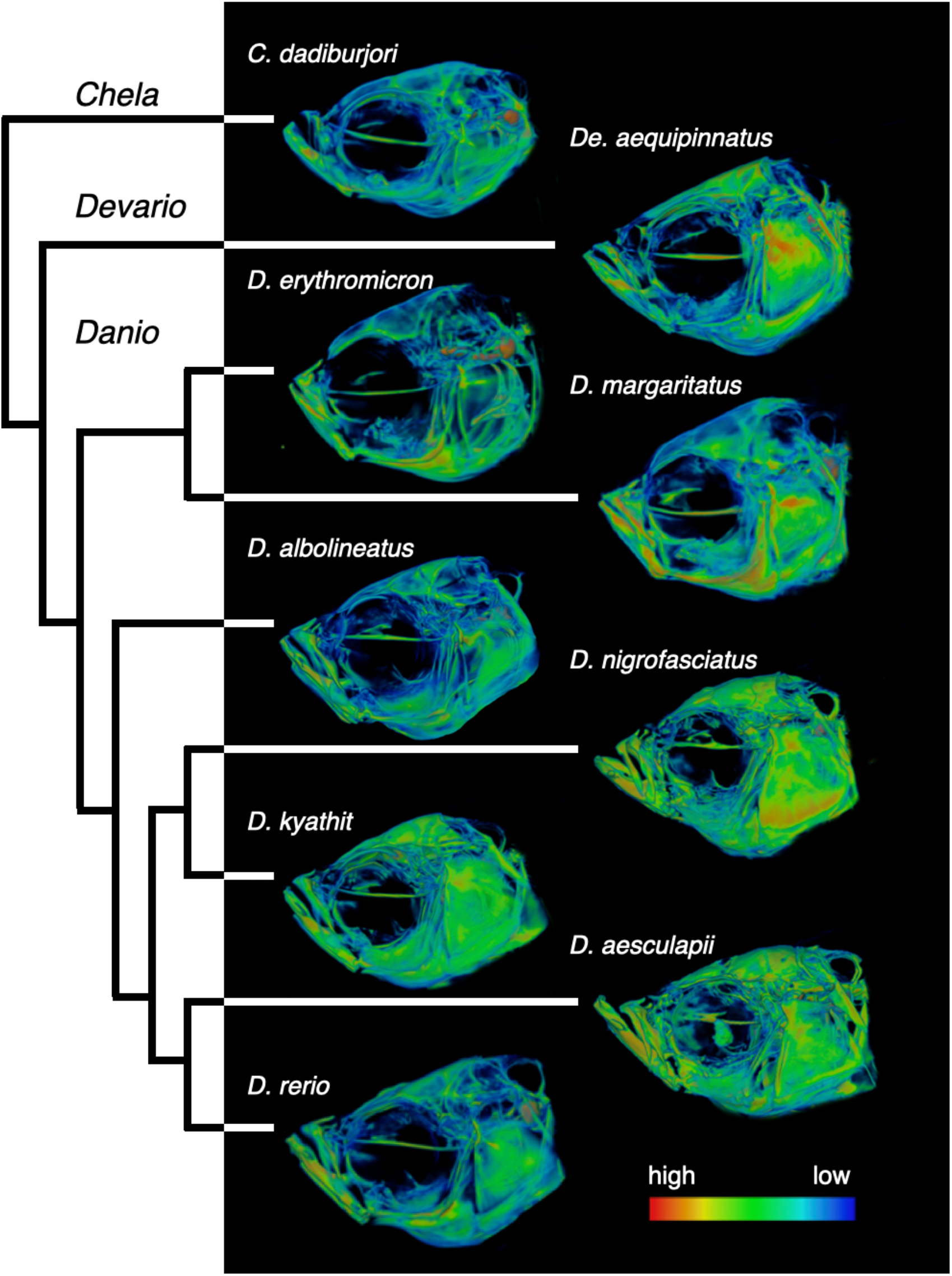
Danioninae show a range of craniofacial morphologies. Left, phylogenetic tree showing relationships between this set of Danioninae species (13). Right, relative density renderings of craniofacial skeletons from nine different Danioninae species. Warmer colors indicate regions of higher relative density. Images and phylogeny are not to scale.

Geometric morphometrics defined the morphospace occupied by these skulls (Fig 2). The first principal component (PC1) described 29% of the variation in the dataset and, from a lateral perspective, captured the elongation of the head and the protrusion of the mouth. PC2 described 23% of the variation, and generally captured the aspect ratio of the head. Notably, all the *Danio* species evaluated were relatively clustered within morphospace, while the *Devario* and *Chela* species did not overlap with any *Danio* individuals (Fig 2). PC2 clearly separated *Danio* from non-*Danios* (*Devario* and *Chela*), with the *Danio* species all showing low PC2 scores and overlapping considerably along this axis.

**Figure 2.**
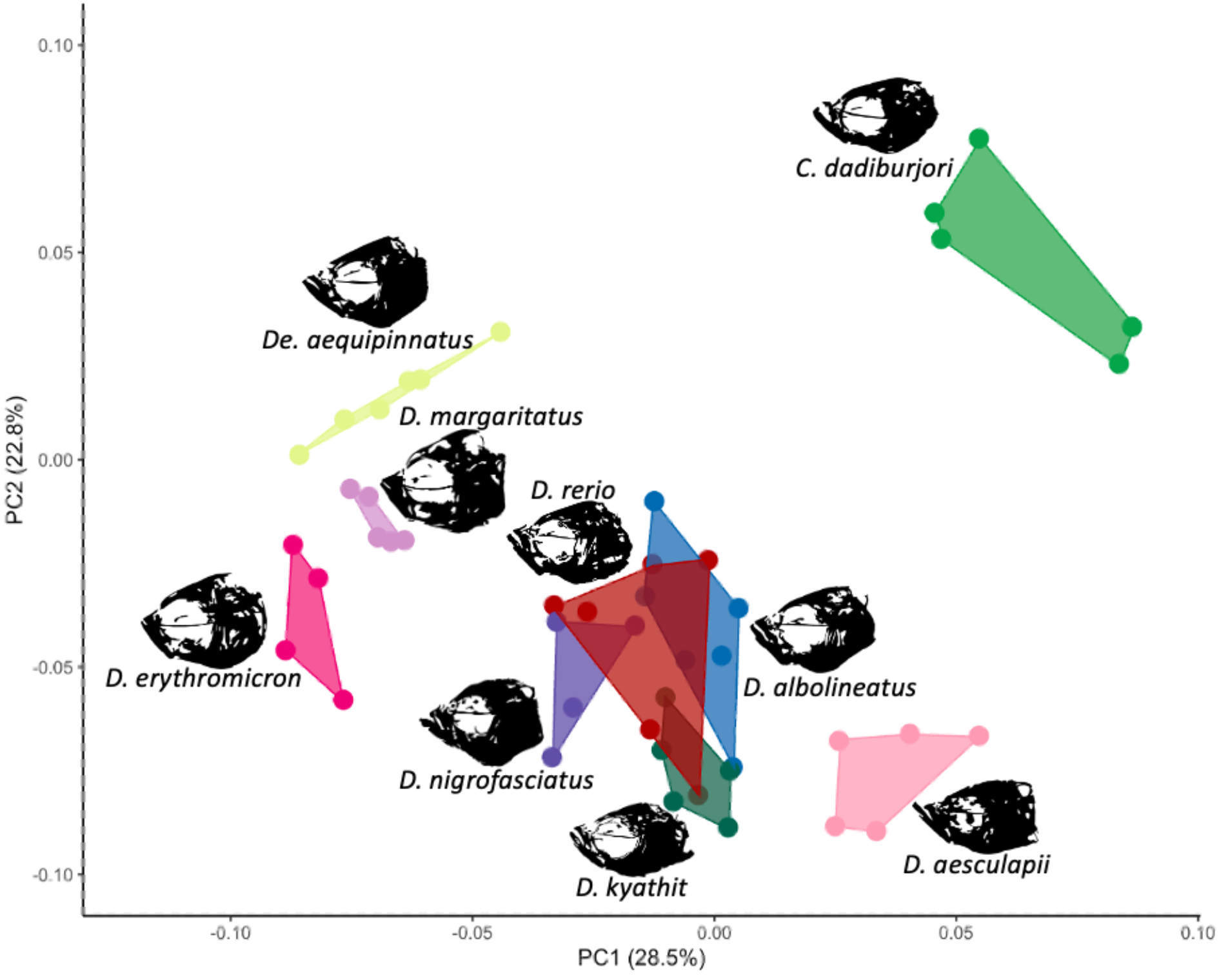
Similarities and differences across Danionin skulls. Morphospace showing the first and second principal components. Multiple individuals were examined for each species and each dot represents a single individual. Polygons represent the area inhabited by all individuals within each species. Shown in black are silhouettes from density renderings representatives of each species.

### Developmental thyroid hormone influences adult skull morphology

To determine the roles that TH plays in shaping *Danio* skulls, we reared *D. rerio* and *D. albolineatus* under altered TH profiles. HypoTH *Danios* showed altered skull shapes and decreased bone density (Fig 3, also see 19, 22, 29). HypoTH conditions induced distinct craniofacial morphologies in both species (Fig 4A), with a distinctly thinner dermatocranium and a somewhat flatter dorsal profile (Fig 3). In a morphospace created by each species reared under different TH conditions, PC2 robustly captured species differences between *D. rerio* and *D. albolineatus*, although it only described 12% of the variation. PC1 (22% of variation) reliably captured the TH status of individuals—whether fish produced TH during development (high PC1) or were HypoTH (low PC1). Notably, HypoTH conditions induced analogous changes in each species.

**Figure 3.**
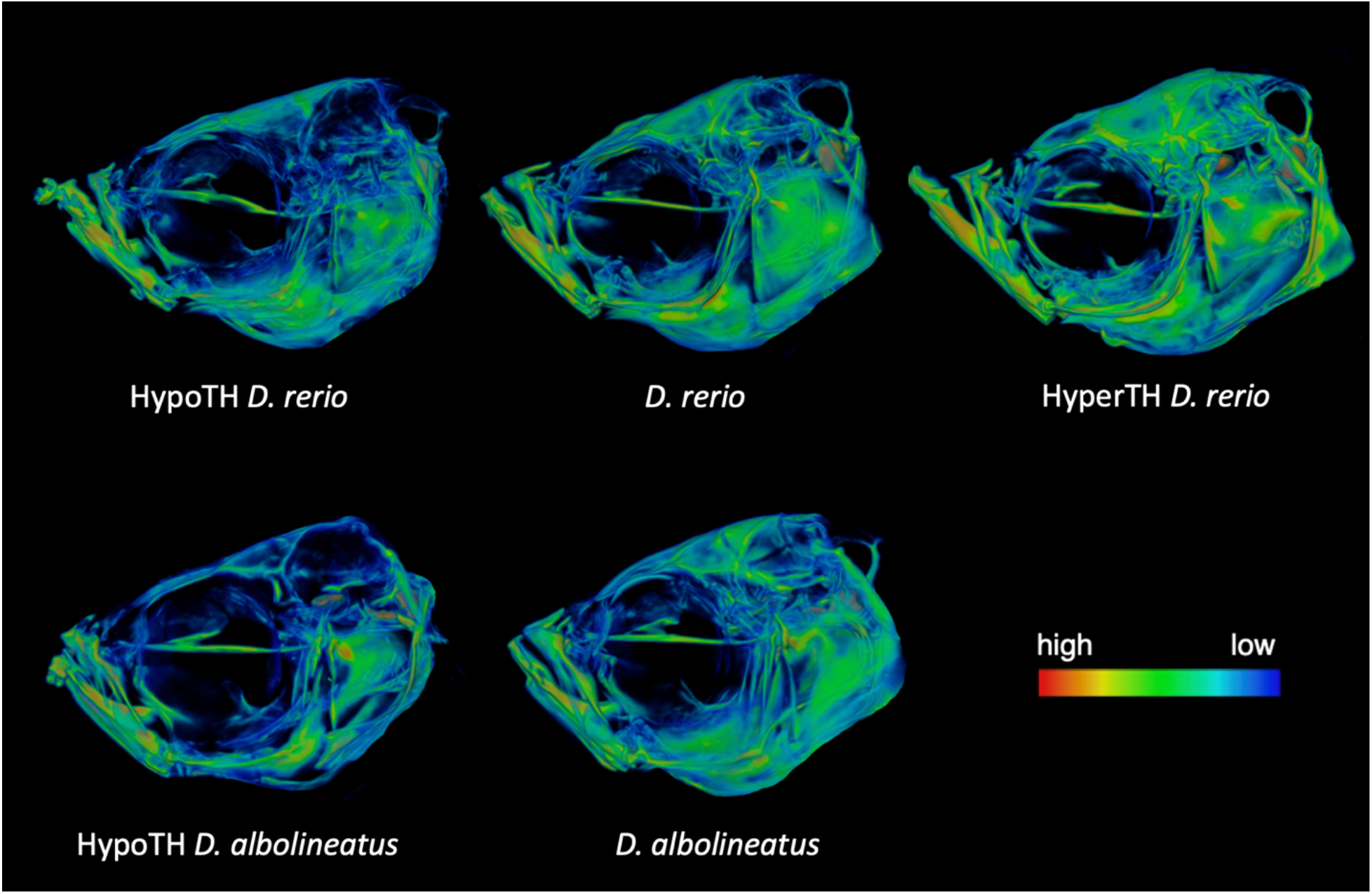
Thyroid hormone regulates craniofacial morphology in two *Danios*. Relative density renderings of hypothyroid (HypoTH), wild-type, and hyperthyroid (HyperTH) *D. rerio* heads. Warmer colors indicate regions of higher relative density.

**Figure 4.**
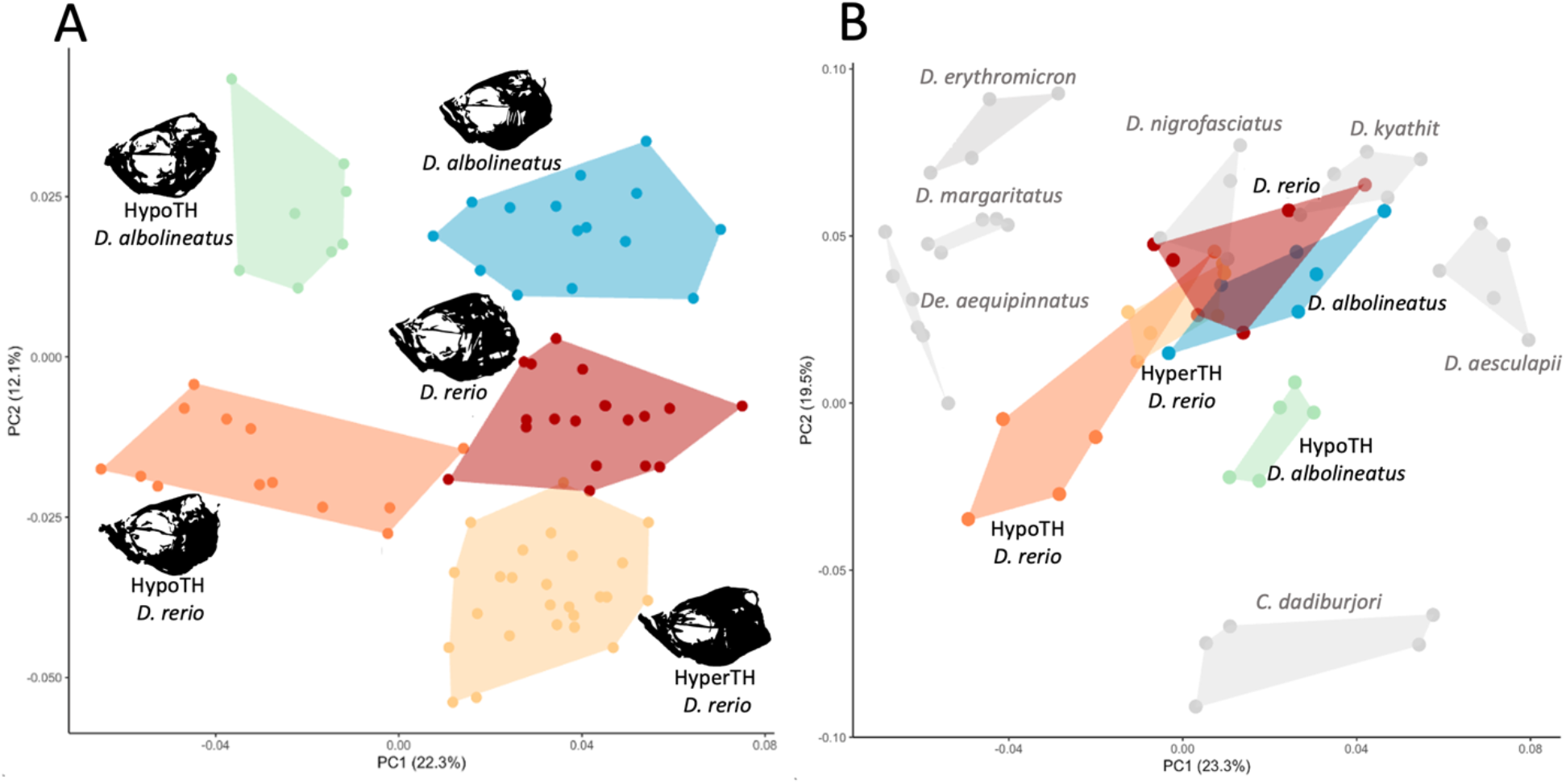
Thyroid hormone availability alters craniofacial phenotypes in evolutionary relevant directions. A, Morphospace defined by *D. rerio* reared under three TH conditions (wild-type, HypoTH and HyperTH) and *D. albolineatus* reared in two TH conditions (wild-type and HypoTH). B, Morphospace defined by all nine species as well as the altered TH conditions. Colors of groups are consistent with those in A; all species shown in Fig 1 are also shown here in grey. In order to have equivalent sample sizes between categories, the best and most representative samples were selected from each group in A and used for the combined PCA shown in B.

HyperTH conditions in *D. rerio* did not induce such marked changes in overall morphology (Fig 4); nonetheless the lower jaw was lengthened by ∼7% in the HyperTH condition (p<<0.001). We note the lower jaws were slightly longer in HypoTH conditions compared to wild-types as well (∼5%; p=0.04), however this was due to the ectopic ossification frequently present at the tips of jaws from HypoTH individuals (visible in Fig 3 and see 29).

### Lack of TH causes *Danio* skulls to more closely resemble other Danioninae

Altering developmental TH concentrations created unique overall craniofacial phenotypes in *D. rerio* and *D. albolineatus*. Putting these TH alteration-induced phenotypes into the context of natural diversity defined a morphospace where PC2 (20% of variation) separated *Danio* and *Devarios* from *Chela* (Fig 4B). Importantly, modified TH conditions caused *D. rerio* and *D. albolineatus* to move out of the morphospace occupied by wild-type *Danio* species in this dataset and closer to the shapes characterizing the *Devario* and *Chela* species evaluated.

## Discussion

Unlike cichlids, which show extraordinary radiations in feeding specializations, Danioninae are typically generalists with multiple species consuming overlapping sources of food (10). Nonetheless, we detected distinct differences in the skull morphologies between different Danioninae species. Most distinct was *C. dadiburjori*, which was the phylogenetic outgroup of the species examined (Fig 1). *Chela* species show certain craniofacial adaptations, including overall elongation, loss of craniofacial barbels and lack of a growth at the tip of the lower jaw (symphyseal knob; see 30). *De. aequipinnatus* also showed a distinct craniofacial morphology. Like other Danioninae, *Devario* species protrude their jaws during feeding, but do so to retain prey rather than to produce suction (4). These species also exhibit premaxilla and skull shapes different from those of *Danio* (4, 19).

The *Danio* species examined in this study all showed similar craniofacial morphologies. Indeed, four of the five crown-group *Danio* (*D. albolineatus, D. nigrofasciatus, D. kyathit* and *D. rerio*) occupied overlapping regions of morphospace (Fig 2). *D. aesculapii*, despite being the sister species to *D. rerio* (13), showed skulls with morphology that did not overlap with that of any other *Danio*. This species notably showed a large lower jaw, reminiscent of the elongated jaws of HyperTH *D. rerio* (19, 29).

TH is required to regulate diverse aspects of zebrafish development, including pigmentation and fin ray morphology (22, 31). The hormone has been shown to influence the development of feeding strategies, and alterations to TH metabolism have been proposed as a mechanism for morphological adaptation of feeding apparatus (4, 19, 22). TH influences the morphology of individual craniofacial elements (21, 29), and here we have shown the hormone to be required for shaping the overall skull in multiple *Danio* species.

Both *D. rerio and D. albolineatus* require TH for the development of normal overall skull morphology. Both species show corresponding morphological shifts when deprived of the hormone during development (Fig 4A). The resulting phenotypes from HypoTH rearing conditions are unlike those seen in skulls of other *Danio* species examined. HypoTH conditions cause skulls to more closely resemble the those of the *Chela* and *Devario* species examined. Changes to developmental TH have the capacity to create unique skull phenotypes that more closely resemble evolved shapes in other Danioninae, adding additional weight to the possibility that shifts in this hormonal axis may be a target for evolutionary morphological change.

## Acknowledgements

Thank you to McMenamin Lab members past and present. Thank you to the lab of David Parichy for generously sharing lines, to BC Core funding and to the members of the BC Animal Care Facility. Funding provided by NSF CAREER 1845513 and R35GM146467 (SKM).

---

**Supplementary Figure 1.**
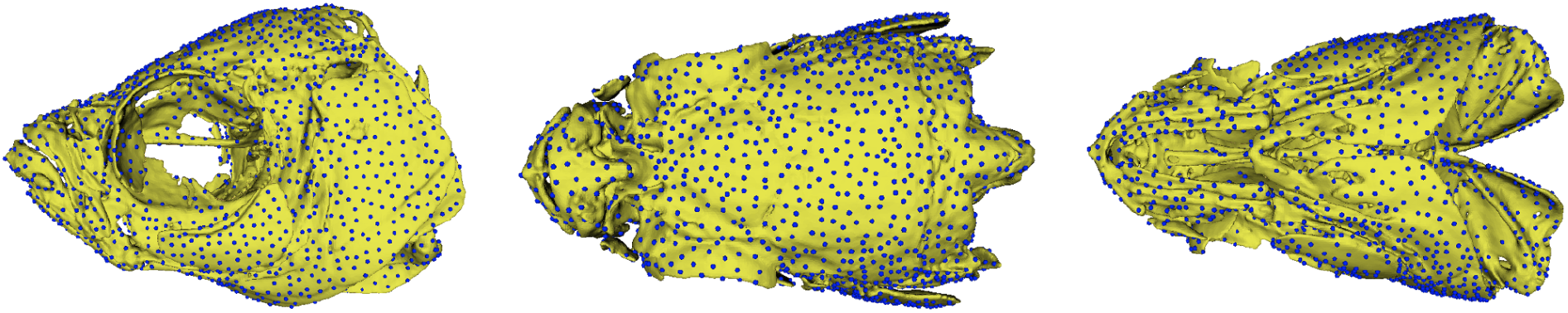
Automatic landmarks to capture 3D shape information. Lateral, dorsal, and ventral views of a 23.0 mm standard length *D. rerio* head with 3000 automatic landmarks placed using ALPACA in 3D Slicer.

